# A ZO-2 scaffolding mechanism regulates the Hippo signalling pathway

**DOI:** 10.1101/2024.10.17.618965

**Authors:** Olivia Xuan Liu, Lester Bocheng Lin, Soumya Bunk, Tiweng Chew, Selwin K. Wu, Fumio Motegi, Boon Chuan Low

**Author notes:** These authors contributed equally to this work (Olivia Xuan Liu, Lester Bocheng Lin, Soumya Bunk).

## Abstract

Inhibition of proliferation upon contact is a critical cell density control mechanism governed by the Hippo signalling pathway. The biochemical signalling underlying cell density-dependent cues regulating Hippo signalling and its downstream effectors, YAP, remains poorly understood. Here, we reveal that the tight junction protein ZO-2 is required for the contact-mediated inhibition of proliferation. We additionally determined that the well-established molecular players of contact inhibition of proliferation, namely Hippo kinase LATS1 and YAP, are regulated by ZO-2, and that the scaffolding function of ZO-2 promotes the interaction with and phosphorylation of YAP by LATS1. Mechanistically, YAP is phosphorylated when ZO-2 brings LATS1 and YAP together via its SH3 and PDZ domains, respectively, subsequently leading to the cytoplasmic retention and inactivation of YAP. In conclusion, we demonstrate that ZO-2 maintains Hippo signalling pathway activation by promoting the stability of LATS1 to inactivate YAP.

## Introduction

The capacity of cells to regulate their proliferation in response to extracellular cues is essential for tissue organisation. Contact inhibition of proliferation prevents cells from growing after reaching a critical cell density [1, 2]. Loss of contact inhibition of proliferation is a hallmark of many cancer cells[3], which have lost their ability to sense cell density and subsequently exhibit uncontrolled cell proliferation and tissue overgrowth[1, 2]. The Hippo pathway has emerged as a conserved signalling pathway that controls cell proliferation in response to the extracellular environment [4–8]. The Hippo signalling pathway consists of several protein kinases, which regulate the activity of the transcriptional coactivator YAP1 (YAP), and its paralog, WW domain-containing transcription regulator protein 1/Transcriptional coactivator with PDZ-binding motif (TAZ) [5–7]. YAP and TAZ bind to transcriptional enhancer associate domain (TEAD) transcription factors and coactivate the expression of proliferation-promoting genes [9–13]. The Serine/threonine-protein kinase 4/Mammalian STE20-like protein kinase 1 (MST1) and Serine/threonine-protein kinase 3/Mammalian STE20-like protein kinase 2 (MST2) kinases trigger phosphorylation and activation of the LATS kinases [14–17], which in turn phosphorylates and inactivates YAP/TAZ by restricting them to the cytoplasm [8, 10, 18, 19]. Hence, the activity of YAP/TAZ is primarily regulated by their subcellular distribution: YAP/TAZ in the cytoplasm are inactive, whereas those in the nucleus are active and stimulate cell proliferation. The subcellular localisation of YAP/TAZ is affected by cell density; YAP/TAZ predominantly accumulates in the nucleus at low cell density and in the cytoplasm at high cell density [8, 20, 21]. As cells grow to confluence, the Hippo kinases are activated and increase the level of phosphorylated YAP in the cytoplasm [8]. These findings suggest that the Hippo pathway plays a crucial role in contact inhibition of proliferation. However, the molecular mechanisms for how cell density is sensed and the signals controlling the subcellular distribution of YAP/TAZ are not well understood.

The distribution of YAP and TAZ can be regulated by multiple pathways, such as cell surface receptor complexes, apical-basal cell polarity, the actin cytoskeleton[22], and cell-extracellular matrix adhesions [5–7, 23]. Because contact inhibition of proliferation relies on cell-cell communication, cell surface adhesion receptors have been attractive key regulators of YAP/TAZ distribution [2, 4, 20, 24]. Of the several classes of cell adhesion receptors, much attention has focused on the protein complex comprising E-cadherin and α/β-catenin[25]. Disrupting functional E-cadherin complexes by the removal of extracellular calcium [20, 26], the addition of anti-E-cadherin antibodies [27], or depletion of β-catenin [21] results in nuclear retention of YAP and TAZ at high cell density and subsequent loss of contact inhibition of proliferation. However, a mutant E-cadherin, which comprises a non-functional extracellular domain but retains a normal membrane-tethered cytoplasmic domain with the ability to bind to α/β-catenins, restored contact inhibition of proliferation [28], suggesting that E-cadherin-based cell-cell junction *per se* is dispensable for contact inhibition of proliferation. Thus, here we focus on the role of Tight junction protein ZO-2 (ZO-2) in contact inhibition of proliferation through regulating Serine/threonine-protein kinase LATS1/ (LATS1) and YAP/TAZ.

ZO-2 is part of the tight junction complex. Tight junctions function as permeability barriers for tissue[29] and play critical roles in many signalling pathways, including the Hippo pathway [30–32]. The Hippo kinases, MST1, MST2, LATS1, and YAP/TAZ accumulate at tight junctions as cell-cell junctions mature[33]. Similarly, ZO-2 also exhibits a dual localisation pattern; like YAP/TAZ, ZO-2 accumulates predominantly in the nucleus at low cell density, whereas ZO-2 localises in the cytoplasm predominantly at high cell density[34]. Although, ZO-2 have been implicated in Hippo signalling[34–38], the biochemical mechanism coupling ZO-2 to Hippo signalling remains elusive. Here, we have identified a biochemical scaffolding mechanism of ZO-2 in regulating Hippo - YAP signalling by affecting LATS1 stability.

## Results & Discussion

### ZO-2 regulates YAP localisation in confluent monolayers

Epithelial cells experience physical constraints from their neighbours as cells proliferate in a confined space, strengthening cell-cell junctions[45], reorganising cytoskeletons and establishing apical-basal polarity[46]. To determine the contribution of external physical constraints on the translocation of YAP from the nucleus to the cytoplasm as cells grow to confluence, we prepared two conditions of MDCK cells at high cell density (Figure 1A). In one condition, cells reach confluency over 72 hours, permitting them to fully polarise gp135/podocalyxin at the apical cortex and E-cadherin at the lateral cell-cell junctions[47] (Figure 1, A-C).

**Figure 1:**
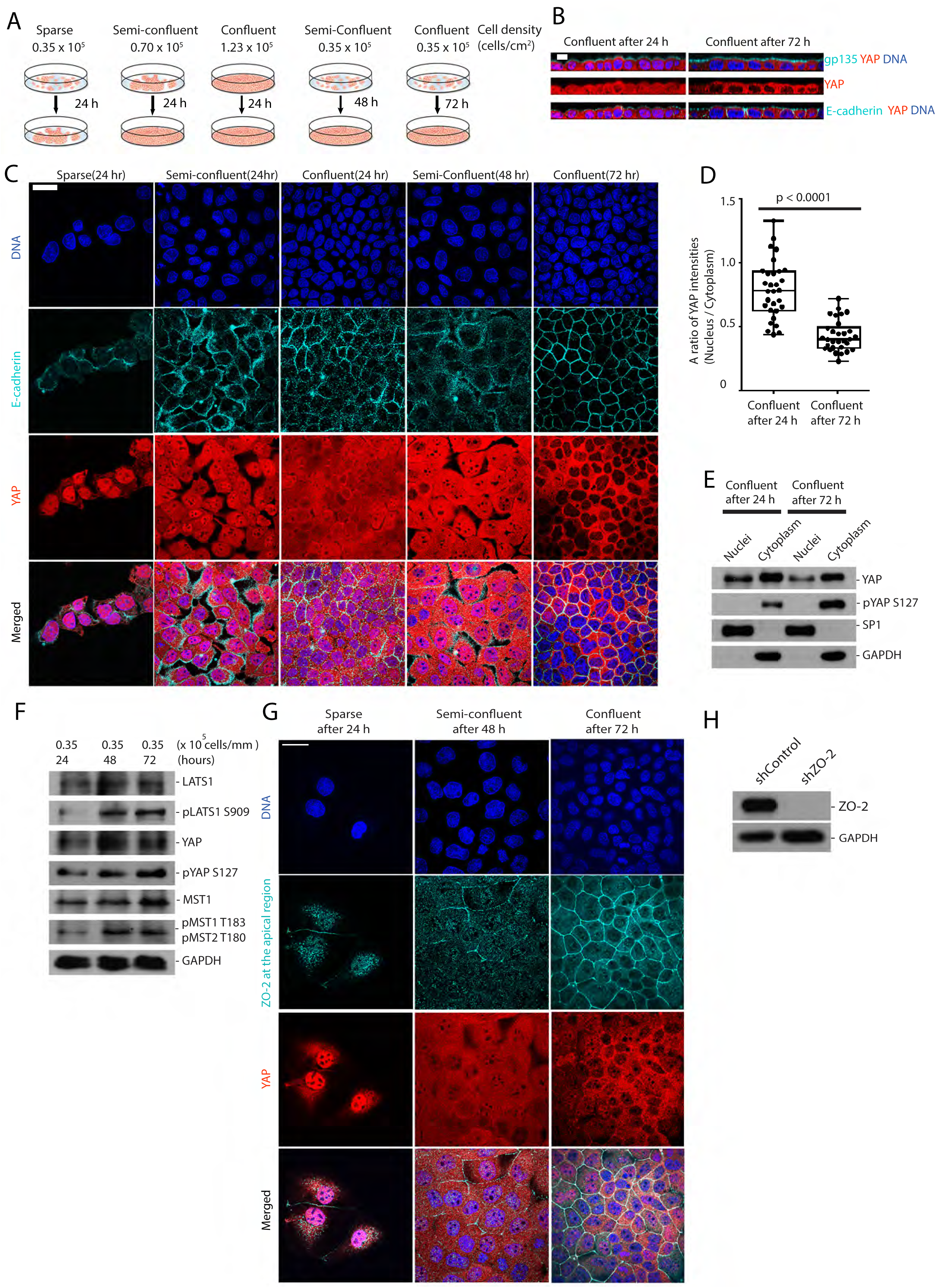
Proper epithelial junction maturation in confluent monolayer is required for YAP nuclear exclusion. A) Preparation of MDCK cells under different cell densities, as indicated. B) Representative cross-sectional views of DAPI, E-cadherin, YAP, and gp135 at different cell densities. C) Images of DAPI, E-cadherin, and YAP immunostained cells seeded at different densities showing a sparse, semi-confluent, or confluent state 24, 48 and 72 hours after cell seeding. D)Quantification of the nuclear to cytoplasmic ratio of YAP intensities in confluent monolayer formed after 24 or 72 hours. D) Representative immunoblots for YAP and S127-phosphorylated YAP (pYAP S127) in the nuclei and the cytoplasmic fraction. SP1 and GAPDH were used as nuclear and cytoplasmic fractions markers, respectively. E) Representative immunoblots for LATS1, S909-phosphorylated LATS1 (pLATS1 S909), YAP, S127-phosphorylated YAP (pYAP S127), MST1, T183/T180-phosphorylated MST1/2 (pMST1 T183/pMST2 T180), and GAPDH. Cell culture conditions of the initial cell seeding densities and elapsed time after seeding are indicated above the immunoblot. F) Representative DAPI, YAP, and ZO-2 images under the indicated cell-density conditions. G) Representative immunoblots of ZO-2 and GAPDH for ZO-2 depleted cells and control. All data are from at least three independent experiments. Scale bar, 25 μm.

In the second condition, cells grew to confluence within 24 hours, which only enabled a weaker accumulation of gp135 at the apical cortex and a weaker accumulation of E-cadherin at the cell-cell junctions (Figure 1, A-C). As expected, YAP localised to nuclei in cells grown under sparse cell density conditions (Figure 1C). At high cell density, YAP was predominantly localised to the cytoplasm in the 72-hour confluent monolayer but was seen both in the cytoplasm and the nucleus in the 24-hour confluent monolayer (Figure 1, B-D). Consistent with our immunofluorescence data, immunoblots show that the amount of phosphorylated YAP (at the S127 residue) and LATS1(at the S909 residue) in the cytoplasm were higher in monolayer formed after 72 hours of seeding (Figure 1, E and F). By contrast, the phosphorylation of MST1 and MST2 (at the T183 and T180 residues, respectively) remains the same in both culturing conditions (Figure 1F). These results confirm that the establishment of apicobasal polarity and epithelial cell-cell junctions promote Hippo signalling pathway activation.

We hypothesised that the tight junction may regulate the Hippo signalling pathway. ZO-2 is an appealing candidate among the tight junction components because, similar to YAP, ZO-2 predominantly accumulates in the nucleus at low cell density, in the cytoplasm, and in the tight junctions at high cell density (Figure 1G). Knockdown of ZO-2 using short-hairpin RNA (shRNA) did not affect YAP localization at low cell density conditions (Figures 1H, 2A and B). The 72 hours confluent monolayer depleted of ZO-2 localizes a substantial amount of YAP in the nuclei (Figure 2, C and D). In contrast, the junctional enrichment of ZO-1 (Figure 2C) and apical enrichment of gp135 is unaffected (Figure 2E). Additionally, ZO-2 knockdown did not have obvious effects on the phosphorylation levels of MST1 and MST2 kinases. Nevertheless, YAP phosphorylation at the S127 residue is reduced (Figure 2F).

**Figure 2:**
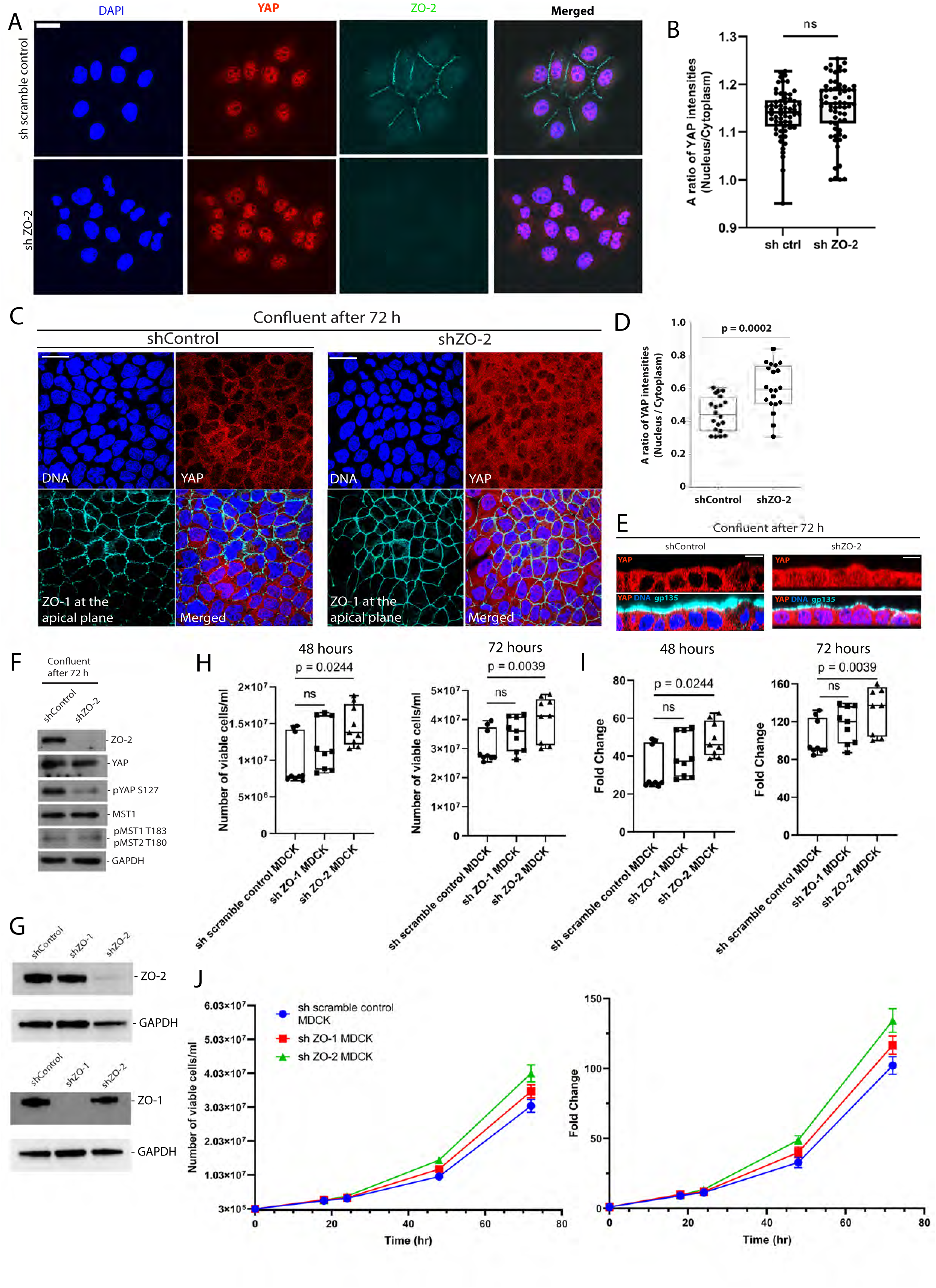
ZO-2 regulates YAP localisation in confluent monolayers. A) ZO-2 is not essential for YAP to enter the nucleus at low cell density. Representative immunostaining of DAPI, ZO-2 and YAP in control and ZO-2 knockdown (KD) under low cell density conditions. B) Quantification of the nuclear to cytoplasmic ratio of YAP intensity at sparse cell density in control and ZO-2 knockdown conditions as shown in (A). Data in box-and-whisker plots show median (midline), 25th to 75th percentiles (box), and minimum and maximum (whiskers) from n>60 cells for each condition, analysed with the Mann-Whitney U test. C) Representative images of DAPI, YAP, and ZO-1 immunostaining of control and ZO-2 depleted cells at confluent cell density 72 hours after seeding. (ZO-1 is at the apical plane, whereas YAP and DAPI are at the lateral plane). D) Quantifying the YAP nuclear to cytoplasmic ratio in cells shown in (C). Data in box-and-whisker plots show median (midline), 25th to 75th percentiles (box), and minimum and maximum (whiskers) from n = 20 cells for each condition, analysed with Student’s t-test. E) Representative cross-sectional views of YAP, DAPI, and gp135 immunostaining in control and ZO-2 depleted cells at confluent cell density 72 hours after seeding. F) Representative immunoblots for ZO-2, YAP, S127-phosphorylated YAP (pYAP S127), MST1, T183/T180-phosphorylated MST1/2(pMST1 T183 / pMST2 T180), and GAPDH from control and ZO-2 depleted confluent monolayer. Data is representative of 5 independent experiments. G) Representative immunoblots for ZO-2 and ZO-1 knockdown in MDCK cells. H) ZO-2 knockdown in confluent monolayer increases the cell number compared to the corresponding controls in the 48 and 72-hour time points. Box-and-whisker plots show median (midline), 25th to 75th percentiles (box), and minimum and maximum (whiskers). All data are from 3 independent experiments, with three technical replicates for each condition (n = 9). Data was analysed with the Mann-Whitney U test. I) ZO-2 knockdown in confluent monolayer increases the fold change of cell proliferation in the 48 and 72-hour time points when compared to the corresponding control. Box-and-whisker plots show median (midline), 25th to 75th percentiles (box), and minimum and maximum (whiskers). All data is from 3 independent experiments, with three technical replicates for each condition (n = 9). Data was analysed with the Mann-Whitney U test. J) ZO-2 knockdown promotes cell proliferation compared to ZO-1 knockdown and control. Proliferation curve of MDCK cells over 72 h. Error bars represent mean ± s.e.m from 3 independent experiments, each with three technical replicates (n=9). Both graphs represent the number of viable cells/ml and fold change relative to seeding cell density. All data are from at least three independent experiments. Scale bar, 25 μm.

A key functional role of Hippo signalling at tight junctions, likely, is to coordinate contact inhibition of proliferation. Thus, we assess ZO-2 role in contact inhibition of proliferation. We quantified MDCK cell proliferation when ZO-2 is depleted (Figure 2G). ZO-2-depleted cells exhibited higher proliferation rates in 48- and 72-hour intervals than cells expressing shRNA scramble control and those depleted of ZO-1 (Figure 2, H-J). Thus, the loss of tight-junction component ZO-2 in contact-inhibited monolayer relieves the proliferation inhibition.

### ZO-2 is required for LATS1 stability

We then investigated if ZO-2 regulates LATS1, the upstream inactivator of YAP. We observed active phospho-LATS1 (S909) concentrating at cell-cell junctions, whereas total LATS1 was distributed throughout the cytoplasm and cell-cell junctions (Figure 3A). In ZO-2-depleted cells, immunofluorescence revealed reduced total and active LATS1 levels (Figure 3A).

**Figure 3:**
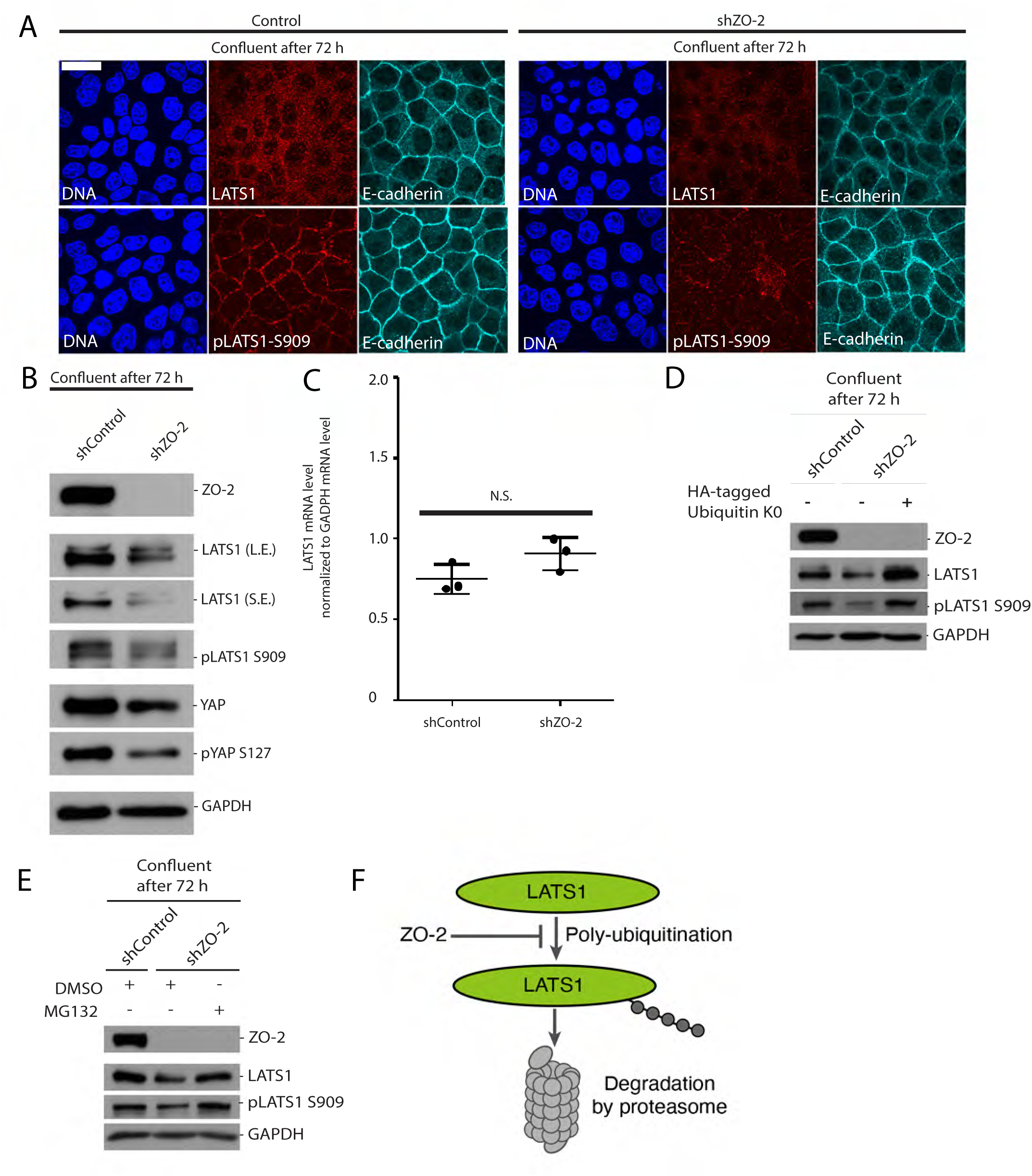
ZO-2 is required for LATS1 stability. A) Representative images of S909-phosphorylated LATS1 (pLATS1 S909) and LATS1, immunostaining in control and ZO-2-depleted cells. E-cadherin staining represents the cell-cell junctions. B) Representative immunoblots for ZO-2, endogenous LATS1 (more prolonged exposure: LE, shorter exposure: SE), S909-phosphorylated LATS1 (pLATS1 S909), YAP, S127-phosphorylated YAP (pYAP S127), and GAPDH from extracts of control and ZO-2 depleted cells. C) LATS1 mRNA level is unaffected by ZO-2 knockdown. RT-qPCR of the levels of LATS1 mRNA normalised to GAPDH mRNA. Values are ± s.d. analysed with the Mann-Whitney U test. D) Inhibition of poly-ubiquitination prevents the reduction of LATS1 protein level in ZO-2-depleted cells. Representative immunoblotting images for ZO-2, LATS1, S909-phosphorylated LATS1 (pLATS1 S909) and GAPDH from control cells and ZO-2 depleted cells with or without the expression of HA-ubiquitin KO. E) Inhibiting proteasome activity increases the LATS1 protein level in ZO-2-depleted cells. Representative immunoblotting images for ZO-2, LATS1, and S909-phosphorylated LATS1 (pLATS1 S909) and GAPDH from extracts of control cells and ZO-2 depleted cells with or without MG132 treatment are shown. Data are from at least two independent experiments. F) A diagram depicting the involvement of ZO-2 in affecting the polyubiquitination and degradation of LATS1. All data are from at least three independent experiments. Scale bar, 25 μm.

Consistent with our immunofluorescence, we observed a significant reduction of LATS1 in ZO-2 knockdown cells in immunoblots (Figure 3B). However, qPCR analysis revealed that the LATS1 mRNA level was unaffected by ZO-2 knockdown (Figure 3C). Given that LATS1 can be poly-ubiquitinated by the ubiquitin ligases[48], we consider the possibility that ZO-2 could affect ubiquitination and consequently prevent the degradation of the LATS1 protein. Indeed, the over-expression of a dominant-negative ubiquitin allele, Ubi-K0, and treatment with a proteasome inhibitor MG132, prevents the degradation in LATS1 protein levels in ZO-2-depleted cells (Figure 3, D and E). Thus, our data suggest that ZO-2 is required for LATS1 protein stability in confluent monolayers (Figure 3F).

### ZO-2 scaffolds LATS1 and YAP to form a complex in confluent monolayers

Next, we examined if ZO-2 promotes the interaction between LATS1 and YAP in a confluent monolayer. Immunoprecipitation assays revealed that endogenous ZO-2 was associated with LATS1 and YAP. LATS1, ZO-2, and YAP interactions were enhanced as the cells grew to confluence (Figure 4, A and B). Consistent with the role of ZO-2 in the recruitment of S909-phosphorylated LATS1 to cell-cell junctions, S909-phosphorylated LATS1 was detected in the pull-down of ZO-2 from the lysates of confluent monolayer (Figure 4C). Depletion of ZO-2 significantly reduced the interaction between LATS1 and YAP (Figure 4D). These results suggest that LATS1–ZO-2–YAP forms a complex in confluent monolayers.

**Figure 4:**
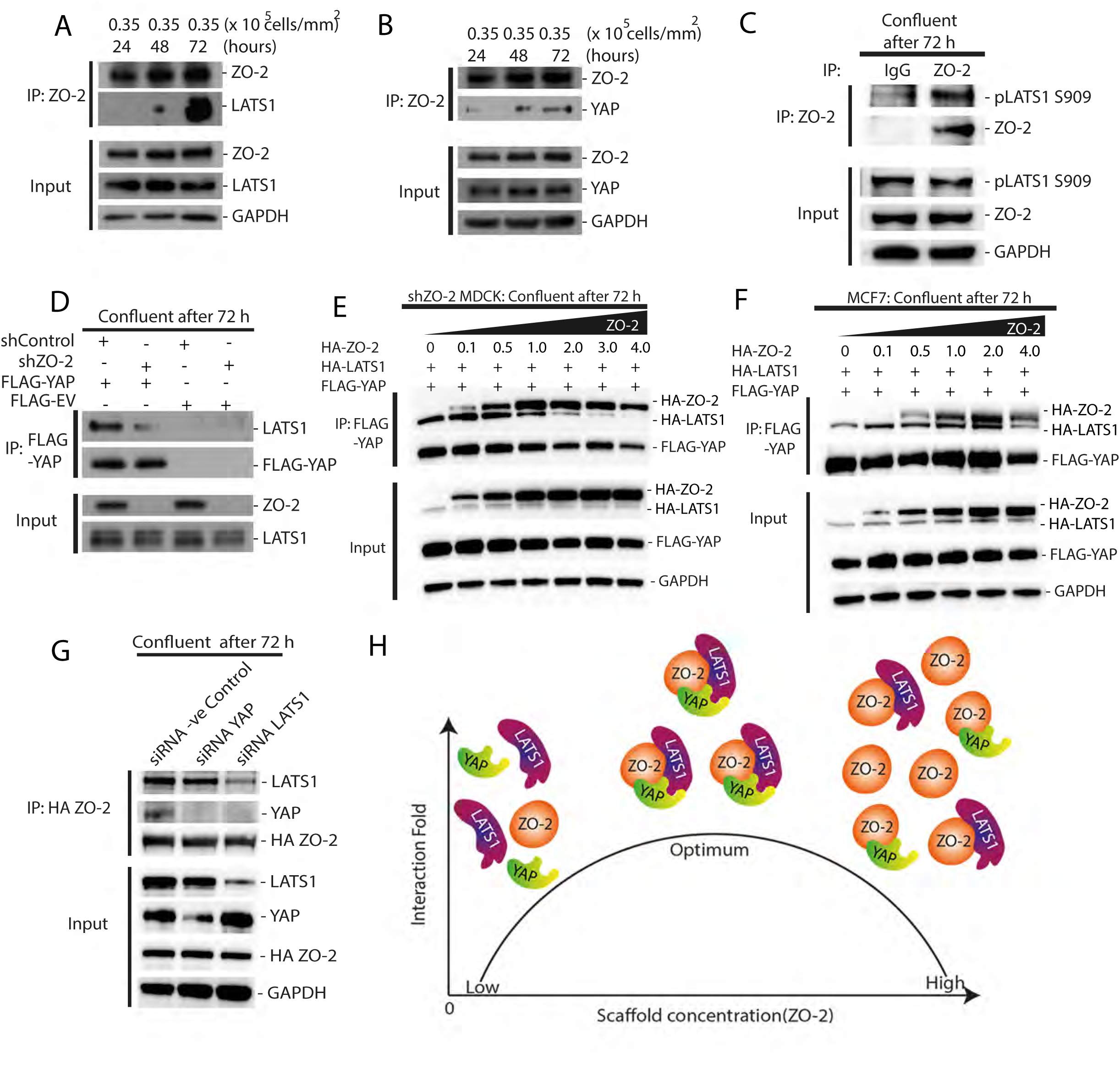
ZO-2 scaffolds LATS1 and YAP to form a complex in confluent monolayers. A) Representative immunoblots of ZO-2 and LATS1. ZO-2 was immunoprecipitated from cells cultured at different densities. ZO-2, LATS1, and GAPDH levels in cells are indicated as inputs. B) Representative immunoblots of ZO-2 and YAP from the ZO-2 immunoprecipitates. ZO-2 was immunoprecipitated from cells cultured at different densities. ZO-2, YAP and GAPDH levels in cells are indicated as inputs C) Representative immunoblots of ZO-2 and pLATS1 S909 in ZO-2 immunoprecipitates. ZO-2 was immunoprecipitated from confluent monolayer of cells 72 hours post-seeding. pLATS1 S909, ZO-2, and GAPDH levels in cells are indicated as inputs. D) ZO-2 promotes LATS1 and YAP interaction at high cell density. Representative immunoblots for LATS1 and FLAG-YAP in FLAG-YAP immunoprecipitates. FLAG-YAP was immunoprecipitated from control and ZO-2-depleted confluent monolayer. Levels of ZO-2 and LATS1 in cells are indicated as inputs. E) and F) ZO-2 scaffolds LATS1 and YAP interaction. Representative immunoblots for HA-LATS1, HA-ZO-2, and FLAG-YAP in FLAG-YAP immunoprecipitates. FLAG-YAP was immunoprecipitated from confluent monolayers expressing a range of ZO-2 levels. HA-ZO-2 and HA-LATS1 levels in cells are indicated as inputs in sh ZO-2 MDCK cells (E) and MCF7 cells (F), respectively. F) ZO-2, YAP and LATS1 interact with each other. YAP and ZO-2 interaction decreases in LATS1 knockdown. However, LATS1 and ZO-2 interaction marginally decreased in the YAP knockdown condition in the MDCK confluent monolayer. G) Diagram of possible scaffolding and titration effects of ZO-2 on LATS1 and YAP. ZO-2 binds to LATS1 and YAP, thus bringing them closer together and facilitating their interactions. Increasing ZO-2 levels can promote the interaction between LATS1 and YAP (optimum levels of ZO-2) until the concentrations of LATS1 and YAP are exceeded by ZO-2. Under conditions of high levels of ZO-2, ZO-2 titrates LATS1 and YAP into separate complexes, thus inhibiting their interactions. According to this model, overexpression of ZO-2 can exert either stimulatory or inhibitory effects on YAP and LATS1 interactions, depending on the levels of ZO-2 relative to YAP and LATS1 abundance. All data are from at least three independent experiments.

Furthermore, we tested whether ZO-2 has a concentration-dependent scaffolding effect[49–51] on the interaction between LATS1 and YAP. In confluent monolayers of both MDCK and MCF7 cells, a fixed amount of HA-LATS1 and FLAG-YAP was transfected together with a different amount of HA-ZO-2 for coimmunoprecipitation. Increasing levels of HA-ZO-2 led to a corresponding increase in the amount of LATS1 co-immunoprecipitated with FLAG-YAP, which reached a maximal level before gradually reducing as HA-ZO-2 concentration further increases (Figure 4, E and F). Based on our data (Figure 4, E and F), we propose that when ZO-2 was highly saturated, a ZO-2/YAP complex or LATS-1–ZO-2 complex was predominantly observed at the expense of a LATS1–ZO-2–YAP complex (Figure 4H). These results imply that an optimal level of ZO-2 is required to form a LATS1–ZO-2–YAP complex efficiently. We further show that while ZO-2 depletions reduced LATS1 and YAP interaction and depletion of LATS1 reduced ZO-2 and YAP interactions, YAP is dispensable for LATS1 and ZO-2 interaction. These results indicate that ZO-2 and LATS1 are required for ZO-2/LATS1/YAP to interact, whereas YAP, a downstream transducer, is not essential for the complex to form (Figure 4G). Thus, reinforcing ZO-2 role as a scaffold for LATS1 and YAP at tight junctions (Figure 4H).

### ZO-2 interacts with LATS1 via its SH3 domain

We then assessed the region of ZO-2 that is associated with LATS1. A truncated version of HA tagged ZO-2, where the SH3 domain was deleted (HA-ZO-2 ΔSH3), was unable to associate with LATS1 as compared to HA-ZO-2(PDZs+SH3) and full-length HA-ZO-2 (Figure 5, A and B). Thus, suggesting that ZO-2 SH3 domain is required to bind LATS1. Additionally, LATS1 was detected in FLAG-YAP immunoprecipitates from the extract of cells depleted of endogenous ZO-2 and expressed full-length ZO-2 but not ZO-2[ΔSH3] (Figure 5C), suggesting that an interaction between the SH3 domain of ZO-2 and LATS1 is essential for LATS1 binding to YAP.

**Figure 5:**
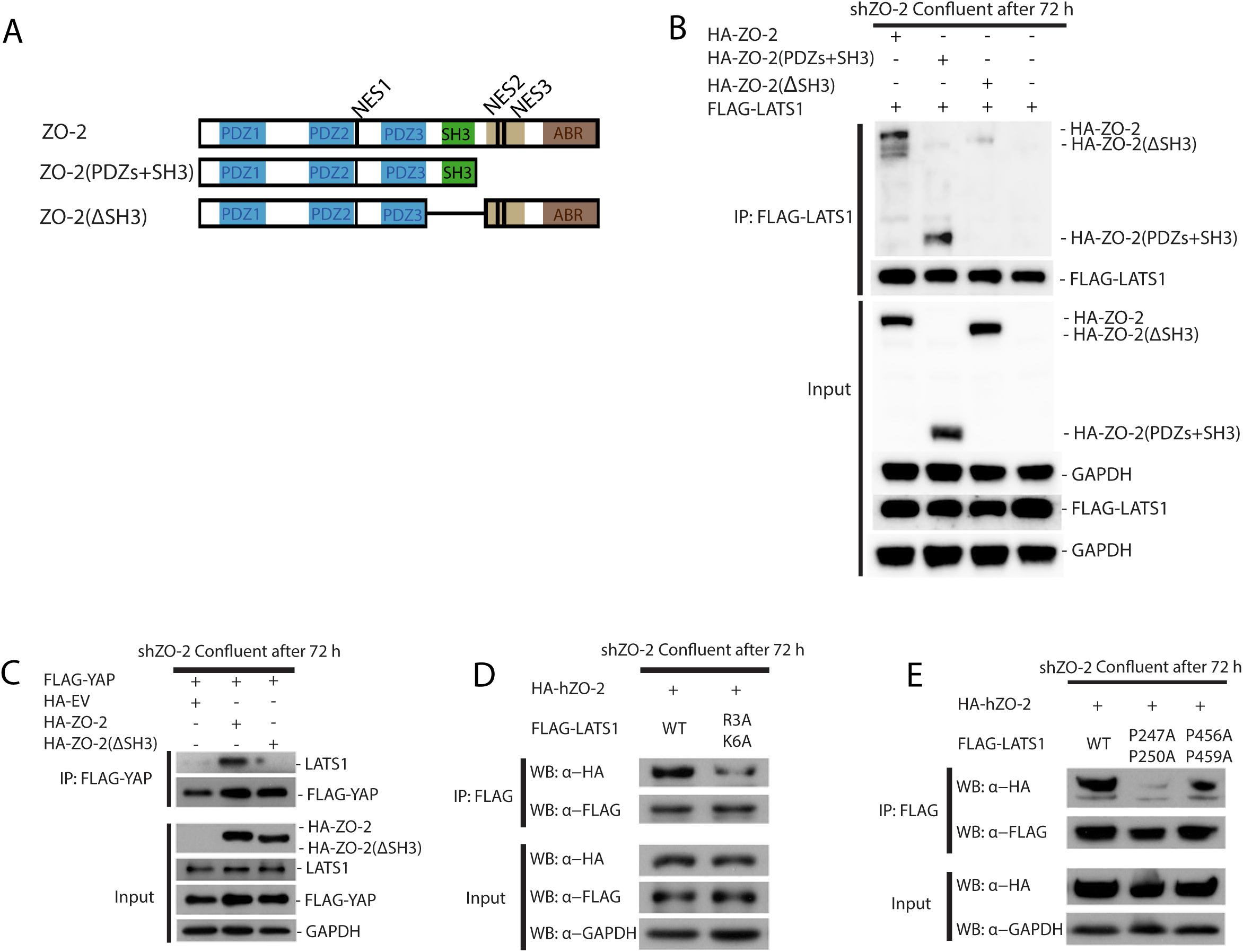
ZO-2 interacts with LATS1 via its SH3 domain. A) Diagram of HA-tagged ZO-2 fragments(PDZ: Post synaptic density protein (PSD95), Drosophila disc large tumor suppressor (Dlg1), and Zonula occludens-1 protein (ZO-1); SH3: Src Homology 3; NES: Nuclear Export Signal; ABR: Actin Binding Region). B) ZO-2 associates with LATS1 via its SH3 domain. Representative immunoblots of HA-tagged full-length ZO-2, ZO-2 deletion mutants and FLAG-LATS1 from FLAG-LATS1 immuno-precipitates. FLAG-LATS1 was immunoprecipitated from sh ZO-2 MDCK confluent monolayers expressing full-length ZO-2 or various ZO-2 deletion mutants. Levels of HA-ZO-2, along with the deletion mutants FLAG-LATS1 and GAPDH, are indicated as inputs. C) The interaction between LATS1 and YAP requires the SH3 domain of ZO-2. Representative immunoblots for LATS1 and FLAG-YAP in the FLAG-YAP immunoprecipitates. FLAG-YAP was immunoprecipitated from ZO-2-depleted cells expressing HA-Empty Vector (EV), HA-ZO-2, or HA-ZO-2(ΔSH3). Levels of HA-ZO-2, HA-ZO-2(ΔSH3), LATS1, FLAG-YAP, and GAPDH are indicated as inputs. D) and E) LATS1 interacts with ZO-2 via its SH3 domain-binding motifs. Alanine scanning mutations were introduced to the SH3-domain-binding motifs of LATS1 (R3A K6A, P247A P250A, P456A P459A). Representative immunoblots for HA-ZO-2, FLAG-LATS1, and GAPDH in FLAG-LATS1 immunoprecipitates are indicated. FLAG-LATS1 were immunoprecipitated from confluent monolayers. Levels of HA-ZO-2, FLAG-LATS1, and GAPDH are indicated as inputs. All data are from at least three independent experiments.

Furthermore, by introducing alanine scanning mutations to the SH3-domain-binding motifs of LATS1 (R3A K6A, P247A P250A, P456A P459A) could not bind to ZO-2 (Figure 5, D and E). These results indicate that ZO-2, through its SH3 domain, binds to LATS1, which enhances the interaction between LATS1 and YAP. Thus, we conclude that ZO-2 serves as a scaffold that enables LATS1 to interact with YAP.

Previously [34], it was reported that siRNA knockdown of ZO-2 (∼62% depletion) did not affect YAP localisation in MDCK cells. By contrast, we have achieved ∼90% ZO-2 knockdown, which enables us to uncover the mechanism of ZO-2 in activating Hippo signalling. This is consistent with other studies that demonstrate ZO-2 plays a role in Hippo signalling[35–38]. However, the biochemical mechanisms remain elusive. Here, we reveal that ZO-2 maintains Hippo signalling activation by stabilizing LATS1 and promoting the scaffolding of LATS1 and YAP to form a LATS1–ZO-2–YAP complex. Of note, the SH3 domain of ZO-2 is critical in this scaffolding mechanism.

We anticipate that our findings could have broad implications. ZO-2 was reported to suppress Yap mediated hepatocyte to cholangiocyte transdifferentiation in the mouse liver[42]. Thus, it is tempting to speculate that the mechanism of ZO2 scaffolding of Yap and LATS1 at tight junctions could prevent hepatocyte to cholangiocyte transdifferentiation[42]. Further, whether the Hippo signalling activation by the ZO-2 scaffolding mechanism is modulated in other contexts, including embryonic development, aging[44, 52] and diseases such as cancer[3, 53], will be a priority for future research[54, 55].

## Materials and methods

### Plasmids

Plasmids for human LATS1 and MST1 expression were gifts from Dr. Zhou Dawang (Xiamen University of China). The LATS1 and MST1 cDNAs were sub-cloned into a series of pXJ40 vectors between the NotI and KpnI and BamHI and XhoI restriction sites, respectively. Human p2xFlag CMV2-ZO-1, p2xFlag CMV2-ZO-2, and p2xFlag CMV2-YAP1-2γ were gifts from Dr. Marius Sudol (National University of Singapore). The ZO-1, ZO-2, and YAP1-2γ cDNAs were sub-cloned into pXJ40 vectors between the XmaI and KpnI, BamHI and KpnI, and NotI and KpnI restriction sites, respectively. The primers used in this study for subcloning and mutagenesis are shown in Table S1.

### Knockdown of ZO-1, ZO-2, and LATS1 by shRNA

Short-hairpin RNAs (shRNAs) targeting ZO-1, ZO-2, and LATS1 were designed using BLOCK-iT^TM^ RNAi Designer according to the p*Silencer*^TM^ siRNA Expression Vectors Instruction Manual instructions. Two single-stranded shRNA template oligonucleotides were heated at 90 °C for 3 minutes and then annealed at 37 °C for 1-2 hours. The annealed shRNA templates were then ligated into HuSH shRNA GFP cloning vector pGFP-V-RS (TR30007). HuSH 29-mer non-effective scrambled shRNA cassettes and pGFP-V-RS (TR30013), were used as negative controls for pGFP-V-RS. Cells were grown with DMEM medium containing 4 μg/mL puromycin (Thermo Fisher Scientific) to confluence, seeded into each well of a 96-well plate and screened for stable cell lines that consistently exhibited a reduction in the level of target proteins. Immunoblotting analyses were used to quantify protein levels in extracts from cells treated with or without shRNA. The primers used in this study for creating the shRNA constructs are shown in Table S2. The knockdown MDCK cells were maintained in a DMEM medium containing 4 μg/mL puromycin, with a media change after every 48 hours.

### Cell culture and transfection

MDCK cells(MDCK-II (RRID:CVCL_0424)) were obtained from Dr. David M. Bryant (University of California, San Francisco). The cells have not been authenticated in our lab. Cells were grown in Dulbecco’s Modified Eagles medium (DMEM) with high glucose (Hyclone) supplemented with 10 % fetal bovine serum (Gibco) and 0.2 % sodium bicarbonate (Sigma-Aldrich) at 37 °C in a humidified incubator supplied with 5 % of CO2. Cells were transfected with 0.5-5.0 μg plasmids in 125 μL Opti-MEM® medium (Thermo Fisher Scientific) with 7.5 μL of Lipofectamine® 3000 reagent (Invitrogen). For immunoblotting experiments, cells were seeded in a 35 mm glass-bottom high *`μ`*-Dish (ibidi) at three distinct cell densities (0.35 x 10^5^, 0.70 x 10^5^, and 1.23 x 10^5^ cells/cm2) to reach sparse, semi-confluent, and confluent culture conditions, respectively, 24 hours after seeding. Alternatively, cells were seeded at a sparse condition (0.35 x 10^5^ cells/cm2) and grown for 24, 48, and 72 hours to reach sparse, semi-confluent, and confluent conditions, respectively. The number of cells was measured using LUNA^TM^ Automated Cell Counter (Logos Biosystems). For immunofluorescence staining experiments, cells were seeded at three distinct cell densities (0.35 x 10^5^, 0.70 x 10^5^, and 1.23 x 10^5^ cells/cm2) and fixed 24 hours post-seeding. Cells were treated with 10 μM MG132 proteasome inhibitor (Sigma Aldrich) or DMSO for 4 hours before Western Blotting. For the MDCK cell growth curve, the cells were seeded at a density of 3.0 x 10^5^ cells. The cells were grown over 72 hours. After staining with Trypan Blue solution (1:1 ratio), the cells were counted using LUNA-II™ Automated Cell Counter. All cells were tested negative for mycoplasma.

### Cell extract preparation

Transfected cells were harvested, washed twice with PBS, and lysed with RIPA buffer (150 mM of NaCl, 50 mM of Tris-HCl (pH 7.3), 0.25 mM of EDTA, 1 % (w/v) sodium deoxycholate, 1 % (v/v) Triton X-100, 50 mM of sodium fluoride, 1 mM of β-glycerophosphate, 1 mM sodium orthovanadate, and Complete Protease Inhibitor Cocktail Tablets (Roche Applied Science)). Cell lysates were centrifuged at 21,000 x g for 10 minutes to remove insoluble cell debris. The total protein concentration in each cell lysate was measured using the Pierce^TM^ bicinchoninic acid (BCA) Protein Assay Kit (Thermo Fisher Scientific).

### Immunoprecipitation

Cell extracts were centrifuged at 21,000 x *g* for 10 minutes. Supernatants were incubated with primary antibodies overnight at 4°C, then mixed with Protein A/G PLUS-Agarose (Santa Cruz Biotechnology) for 2 hours at 4°C. The agarose beads were washed thrice with RIPA buffer and treated with 6X protein SDS loading dye at 85 °C for 6 minutes. The FLAG fusion protein was immunoprecipitated with an anti-FLAG antibody conjugated with M2 beads (Sigma-Aldrich).

### Immunoblotting

Proteins in the cell extracts and the immunoprecipitates were separated by SDS-PAGE and then transferred onto a PVDF membrane. The PVDF membrane was incubated with blocking buffer (PBS containing 1 % (w/v) BSA and 0.1 % (v/v) Tween-20) overnight at 4 °C. After incubation with primary antibody in blocking buffer, the membrane was washed three times with washing buffer (PBS, 0.1 % (v/v) Tween-20) at room temperature. Horseradish peroxidase-conjugated anti-mouse antibody (GE Healthcare) was used as a secondary antibody and detected by chemiluminescence using the Pierce^TM^ enhanced chemiluminescence Western Blotting Substrate Kit (Thermo Scientific).

### Primary antibodies used for immunoblotting

Rabbit polyclonal anti-ZO-2 antibody (H-110), rabbit polyclonal anti-ZO-1 antibody (H-300), and rabbit polyclonal anti-Sp1 antibody (H-225) were purchased from Santa Cruz Biotechnology. Rabbit polyclonal anti-LATS1 antibody (#9153), rabbit polyclonal anti-phospho-LATS1 (Ser909) antibody (#9157), rabbit polyclonal anti-phospho-YAP (Ser127) antibody (#4911), rabbit polyclonal anti-MST1 antibody (#3682), rabbit polyclonal anti-phospho-MST1 (Thr183) and MST2 (Thr180) antibody (#3681), rabbit polyclonal anti-Histone H3 antibody (#9715), rabbit polyclonal anti-phospho-Histone H3 (Ser10) antibody (#9701) and rabbit monoclonal anti-YAP antibody (#14074) were purchased from Cell Signalling Technology. Rabbit polyclonal anti-FLAG antibody, goat anti-rabbit IgG (A4914), and anti-mouse IgG antibodies conjugated with horseradish peroxidase (A4416) were purchased from Sigma-Aldrich. Rabbit polyclonal anti-HA antibody (#71-5500) was purchased from Invitrogen. Mouse monoclonal anti-GAPDH antibody (AM4300) was purchased from Ambion® Applied Biosystems. Monoclonal mouse anti-E-Cadherin antibody (#610182) was purchased from BD Transduction Laboratories^TM^. Rabbit polyclonal anti-YAP antibody was a gift from Dr. Marius Sudol (National University of Singapore). Mouse monoclonal anti-GAPDH antibody (#437000) was purchased from Thermo Fisher Scientific.

### Reverse transcription PCR (RT-PCR) and Real-time quantitative PCR (RT-qPCR)

Cells were seeded at the sparse condition (0.35 x 10^5^ cells/cm2) and grown for 72 hours. Cells were subsequently lysed with a buffer RLT (Qiagen) containing 1 % (v/v) ß-mercaptoethanol and then homogenised by passing through a blunt 20-gauge needle fitted to an RNase-free syringe. The total RNA in these cell extracts was purified using RNeasy spin columns (Qiagen). Five micrograms of total RNA were used for first-strand cDNA synthesis with SuperScript® III Reverse Transcriptase Kit (Thermo Fisher Scientific). The first-strand cDNA was then incubated with 0.1 M DTT, RNaseOUT^TM^ Recombinant RNase Inhibitor, and SuperScript^TM^ III reverse transcriptase at 25 °C for 5 minutes, followed by 50 °C for 60 minutes, and then at 70°C for 15 minutes. Synthesised cDNAs were used as templates for amplifying target genes in RT-PCR and RT-qPCR experiments. RT-PCR was performed using 100 ng cDNA template, 0.2 mM dNTPs, DyNAzyme II DNA 1 polymerase (Thermo Fisher Scientific), and 10 mM each of the forward and reverse primers. PCR was performed using the following conditions: initial denaturation at 95 *`°`*C for 2 minutes, followed by 35 cycles of denaturation at 95 °C for 45 seconds, primer annealing at 55 °C for 45 seconds, and elongation at 72 °C for 1 minute, and lastly a final elongation at 72 °C for 10 minutes. RT-qPCR was performed using 80-140 ng cDNA, SsoFast^TM^ EvaGreen® Supermix (Bio-Rad), and 500 nM each of the forward and reverse primers. This reaction mixture was then subjected to an initial denaturation at 95 °C for 30 seconds, followed by 50 cycles of reactions, each consisting of denaturation at 95 °C for 5 seconds and combined annealing and extension at 60 °C for 5 seconds. Melt curves were generated at a T_m_ ranging from 65 °C to 95 °C for 5 seconds at each increment of 0.5 °C.

### Immunostaining

Immunostaining experiments were performed with cells grown in a 35 mm high glass bottom *`μ`*-Dish (ibidi). Cells were washed with PBS and fixed with PBS containing 4 % paraformaldehyde for 15 minutes at 37 °C. Fixed cells were washed with PBS and then with PBS containing ammonium chloride. Cells were then permeabilised with PBS containing 0.2 % Triton X-100 for 5 minutes, incubated with blocking buffer (2 % BSA and 7 % FBS in PBS) for 1 hour at room temperature, and incubated with the primary antibodies in blocking buffer overnight at 4 °C. Subsequently, cells were washed five times using PBS buffer containing 0.1% Triton X-100, and then incubated with the Alexa Fluor® conjugated secondary antibodies and Hoechst 33258 in blocking buffer for 1 hour at room temperature[39–43].

### Antibodies for immunofluorescence

Rabbit polyclonal anti-YAP antibody was a gift from Dr. Marius Sudol (National University of Singapore). Rabbit polyclonal anti-phospho-YAP (Ser127) antibody (#4911), rabbit polyclonal anti-LATS1 antibody (#9153), rabbit polyclonal anti-phospho-LATS1 (Ser909) antibody (#9157), and rabbit monoclonal anti-YAP antibody (#14074) were purchased from Cell Signalling Technology. Rabbit polyclonal anti-ZO-2 (H-110) antibody and goat polyclonal anti-ZO-2 (R-19) were purchased from Santa Cruz Biotechnology. Mouse monoclonal anti-gp135 antibody was obtained from the DSHB (DSHB Hybridoma Product 3F2/D8). Mouse monoclonal anti-FLAG antibody (F3165), rat monoclonal anti-Uvomorulin/E-Cadherin antibody (U3254), and Hoechst 33258 were all purchased from Sigma-Aldrich. Mouse monoclonal anti-HA.11 epitope tag antibodies (16B12) were purchased from Covance. Mouse monoclonal anti-ZO-1 antibody (#33-9100), and all the secondary antibodies conjugated with Alexa Fluor® dye (405, 488, 568 and 633) (A31556 (Goat anti-rabbit, 405 nm), A31553 (Goat anti-mouse IgG, 405 nm), A11070 (Goat anti-rabbit, 488 nm), A21202 (Donkey anti-mouse, 488 nm), A11036 (Goat anti-1 rabbit, 568 nm), A21071 (Goat anti-rabbit, 633 nm), A21052 (Goat anti-mouse, 633 nm) were purchased from Invitrogen.

### Imaging and analysis

Cells were observed at 37 °C with a PlanApochromat 100X 1.4 NA oil immersion lens on a Nikon A1R inverted microscope (Nikon) outfitted with 405 nm, 488 nm, 561 nm, and 640 nm lasers (Coherent Sapphire). The images were taken stepwise at 0.1-0.2 μm per step from the focal plane at the bottom of the cell, through the nucleus, to the top of the cells. The z-stacks were merged and analysed using the Volume Viewer function in ImageJ software. The fluorescence intensities of YAP were measured per square μm within the nucleus relative to that in the cytoplasm, and this calculation determines the fold difference in YAP protein levels between the nucleus and the cytoplasm[44]. For structured illumination microscopy, spinning-disc confocal microscopy (Yokogawa) Nikon Eclipse Ti-E inverted microscope with Perfect Focus System, controlled by MetaMorph software (Molecular device) supplemented with a 60x oil 1.3 NA CFI Plan Apo Lambda oil immersion objective and sCMOS camera (Prime 95B, Photometrics) was used.

The ZO-2 knockdown percentage was measured using ImageJ software. The ZO-2 and GAPDH protein band intensity was measured by subtracting the background intensity, and the ratio of ZO-2 to GAPDH band intensity was measured and compared between the ZO-2 knockdown and control conditions.

### Statistical Analysis

All statistical tests were performed using GraphPad Prism 7.0 and 9.0. Results presented in the graphs represent the mean ± standard error of mean (s.e.m.) or standard deviation (s.d.). The D’Agostino-Pearson omnibus test was used for normality testing of the data. The Student’s t-test (two-tailed distribution) or Mann–Whitney U test (two-tailed distribution) was used to calculate *p*-values. The exact sample numerical value and the statistical methods are indicated in the corresponding Figure or Figure legend. All experiments were independently repeated at least three times with similar results, and the representative data are shown. No statistical method was used to predetermine the sample size. The experiments were not randomised. The investigators were not blinded to allocation during experiments or outcome assessment.

## Author contributions

All authors developed the experimental design together and presented ideas. SKW, FM, TWC and BCL guided the study. SKW, OXL and LBL wrote the manuscript with input from all authors. SB edited the manuscript under SKW’s guidance. OXL performed all experiments with guidance from TWC, except for Figures 1, G-J and O-R, Figure 2A, Figure 3, E-H and Figure 4B, which SB did. LBL analysed images and performed all statistical analyses with input from SB and SKW.

## Acknowledgements

This study was supported by the Mechanobiology Institute of Singapore (to BCL), funded through the National Research Foundation and the Ministry of Education Singapore, the Singapore National Research Foundation (NRF_NRFF2012-08 to FM), and the Strategic Japan-Singapore Cooperative Research Program by the Japan Science and Technology Agency and the Singapore Agency for Science, Technology, and Research (1514324022 to FM). We are grateful to Andrew Wong and Marius Sudol, and members of the LBC lab for their helpful comments on the manuscript.

## Data availability

All data and materials supporting the findings of this study are available from the corresponding author upon reasonable request. The ImageJ macro used in this study for quantifying YAP nuclear/cytoplasmic ratio was used in previous studies[44, 56] and is available upon request.

## Conflicts of interest

The authors declare no competing financial interests.

## Abbreviations

YAP: Yes-associated protein
TAZ: Transcriptional co-activator with PDZ-binding motif
MST1/2: Mammalian STE20-like protein kinase ½
ZO-1/2: Zonula Occludens-1/2
LATS1: Large Tumor Suppressor Kinase 1
SH3: Src Homology 3
MDCK: Madin-Darby Canine Kidney

## Supporting information

**Table S1.**
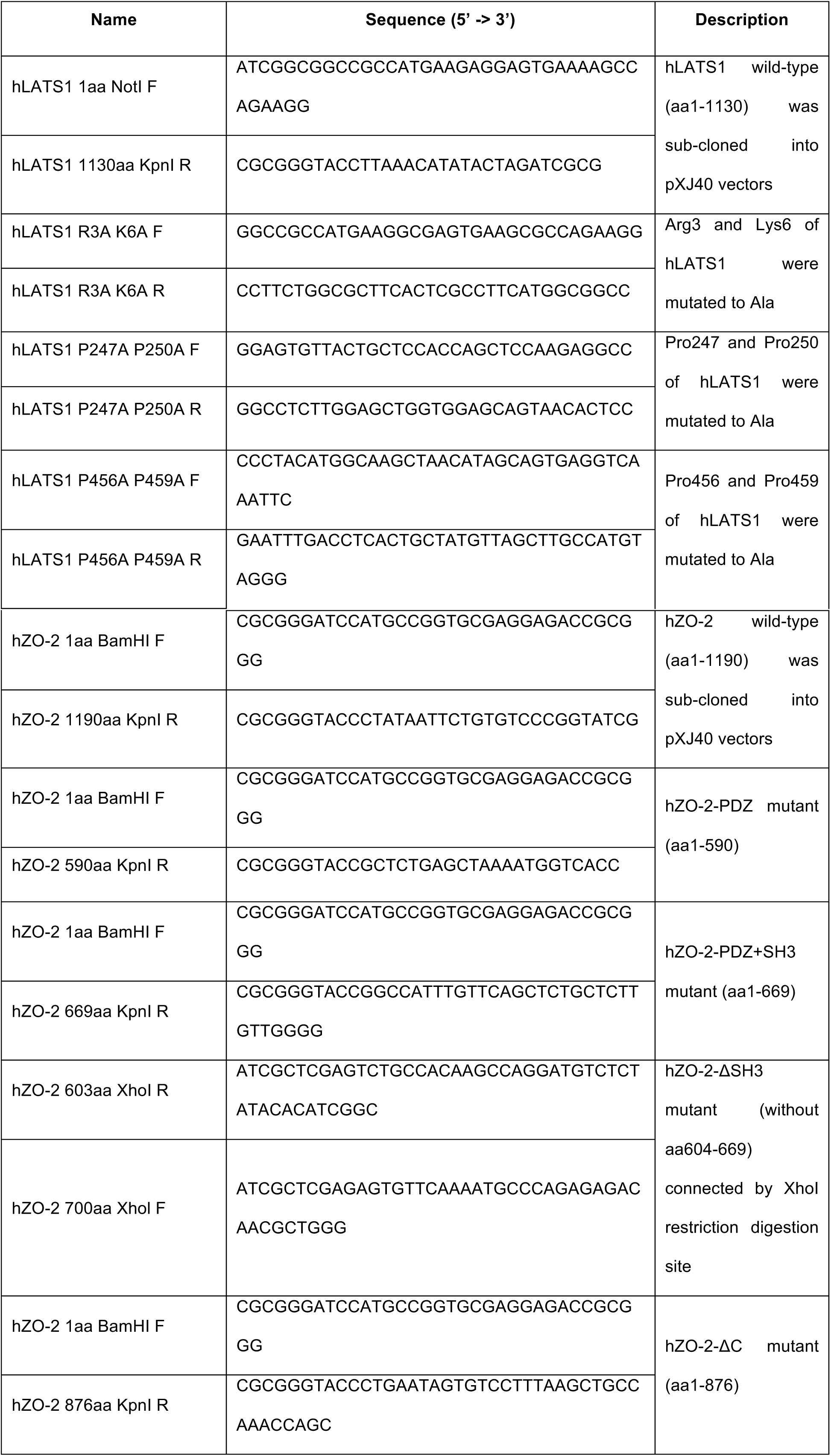
Primers for subcloning and mutagenesis.

**Table S2.**
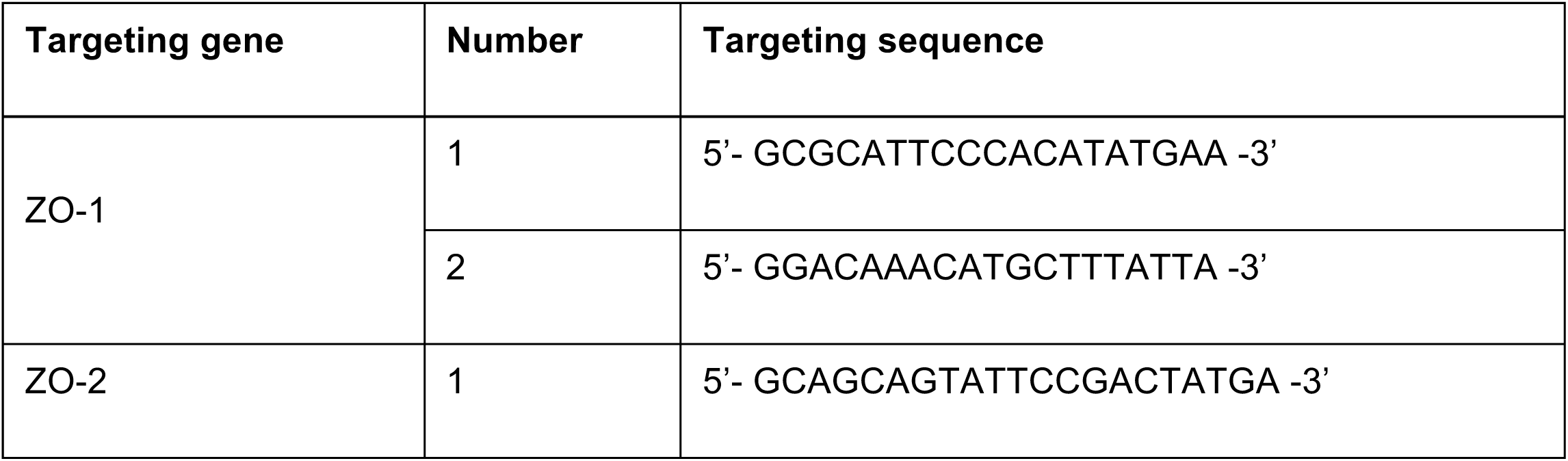
shRNA targeting sequences against LATS1, YAP, ZO-1 and ZO-2 from MDCK cells.

## Notes

https://www.biorxiv.org/content/10.1101/355081v1.full

### Competing Interest Statement

The authors have declared no competing interest.

## References

1. Edgar, B. A. (2006) From cell structure to transcription: Hippo forges a new path, Cell. 124, 267–73.

2. McClatchey, A. I. & Yap, A. S. (2012) Contact inhibition (of proliferation) redux, Curr Opin Cell Biol. 24, 685–94.

3. Wu, S. K., Lagendijk, A. K., Hogan, B. M., Gomez, G. A. & Yap, A. S. (2015) Active contractility at E-cadherin junctions and its implications for cell extrusion in cancer, Cell Cycle. 14, 315–22.

4. Gumbiner, B. M. & Kim, N. G. (2014) The Hippo-YAP signaling pathway and contact inhibition of growth, J Cell Sci. 127, 709–17.

5. Pan, D. (2010) The hippo signaling pathway in development and cancer, Dev Cell. 19, 491–505.

6. Panciera, T., Azzolin, L., Cordenonsi, M. & Piccolo, S. (2017) Mechanobiology of YAP and TAZ in physiology and disease, Nat Rev Mol Cell Biol. 18, 758–770.

7. Yu, F. X., Zhao, B. & Guan, K. L. (2015) Hippo Pathway in Organ Size Control, Tissue Homeostasis, and Cancer, Cell. 163, 811–28.

8. Zhao, B., Wei, X., Li, W., Udan, R. S., Yang, Q., Kim, J., Xie, J., Ikenoue, T., Yu, J., Li, L., Zheng, P., Ye, K., Chinnaiyan, A., Halder, G., Lai, Z. C. & Guan, K. L. (2007) Inactivation of YAP oncoprotein by the Hippo pathway is involved in cell contact inhibition and tissue growth control, Genes Dev. 21, 2747–61.

9. Galli, G. G., Carrara, M., Yuan, W. C., Valdes-Quezada, C., Gurung, B., Pepe-Mooney, B., Zhang, T., Geeven, G., Gray, N. S., de Laat, W., Calogero, R. A. & Camargo, F. D. (2015) YAP Drives Growth by Controlling Transcriptional Pause Release from Dynamic Enhancers, Mol Cell. 60, 328–37.

10. Huang, J., Wu, S., Barrera, J., Matthews, K. & Pan, D. (2005) The Hippo signaling pathway coordinately regulates cell proliferation and apoptosis by inactivating Yorkie, the Drosophila Homolog of YAP, Cell. 122, 421–34.

11. Vassilev, A., Kaneko, K. J., Shu, H., Zhao, Y. & DePamphilis, M. L. (2001) TEAD/TEF transcription factors utilize the activation domain of YAP65, a Src/Yes-associated protein localized in the cytoplasm, Genes Dev. 15, 1229–41.

12. Wu, S., Liu, Y., Zheng, Y., Dong, J. & Pan, D. (2008) The TEAD/TEF family protein Scalloped mediates transcriptional output of the Hippo growth-regulatory pathway, Dev Cell. 14, 388–98.

13. Zanconato, F., Forcato, M., Battilana, G., Azzolin, L., Quaranta, E., Bodega, B., Rosato, A., Bicciato, S., Cordenonsi, M. & Piccolo, S. (2015) Genome-wide association between YAP/TAZ/TEAD and AP-1 at enhancers drives oncogenic growth, Nat Cell Biol. 17, 1218–27.

14. Callus, B. A., Verhagen, A. M. & Vaux, D. L. (2006) Association of mammalian sterile twenty kinases, Mst1 and Mst2, with hSalvador via C-terminal coiled-coil domains, leads to its stabilization and phosphorylation, Febs j. 273, 4264–76.

15. Chan, E. H., Nousiainen, M., Chalamalasetty, R. B., Schäfer, A., Nigg, E. A. & Silljé, H. H. (2005) The Ste20-like kinase Mst2 activates the human large tumor suppressor kinase Lats1, Oncogene. 24, 2076–86.

16. Wei, X., Shimizu, T. & Lai, Z. C. (2007) Mob as tumor suppressor is activated by Hippo kinase for growth inhibition in Drosophila, Embo j. 26, 1772–81.

17. Wu, S., Huang, J., Dong, J. & Pan, D. (2003) hippo encodes a Ste-20 family protein kinase that restricts cell proliferation and promotes apoptosis in conjunction with salvador and warts, Cell. 114, 445–56.

18. Dong, J., Feldmann, G., Huang, J., Wu, S., Zhang, N., Comerford, S. A., Gayyed, M. F., Anders, R. A., Maitra, A. & Pan, D. (2007) Elucidation of a universal size-control mechanism in Drosophila and mammals, Cell. 130, 1120–33.

19. Lei, Q. Y., Zhang, H., Zhao, B., Zha, Z. Y., Bai, F., Pei, X. H., Zhao, S., Xiong, Y. & Guan, K. L. (2008) TAZ promotes cell proliferation and epithelial-mesenchymal transition and is inhibited by the hippo pathway, Mol Cell Biol. 28, 2426–36.

20. Schlegelmilch, K., Mohseni, M., Kirak, O., Pruszak, J., Rodriguez, J. R., Zhou, D., Kreger, B. T., Vasioukhin, V., Avruch, J., Brummelkamp, T. R. & Camargo, F. D. (2011) Yap1 acts downstream of α-catenin to control epidermal proliferation, Cell. 144, 782–95.

21. Kim, N. G., Koh, E., Chen, X. & Gumbiner, B. M. (2011) E-cadherin mediates contact inhibition of proliferation through Hippo signaling-pathway components, Proc Natl Acad Sci U S A. 108, 11930–5.

22. Moore, T., Wu, S. K., Michael, M., Yap, A. S., Gomez, G. A. & Neufeld, Z. (2014) Self-organizing actomyosin patterns on the cell cortex at epithelial cell-cell junctions, Biophys J. 107, 2652–61.

23. Low, B. C., Pan, C. Q., Shivashankar, G. V., Bershadsky, A., Sudol, M. & Sheetz, M. (2014) YAP/TAZ as mechanosensors and mechanotransducers in regulating organ size and tumor growth, FEBS Lett. 588, 2663–70.

24. Silvis, M. R., Kreger, B. T., Lien, W. H., Klezovitch, O., Rudakova, G. M., Camargo, F. D., Lantz, D. M., Seykora, J. T. & Vasioukhin, V. (2011) α-catenin is a tumor suppressor that controls cell accumulation by regulating the localization and activity of the transcriptional coactivator Yap1, Sci Signal. 4, ra33.

25. Wu, S. K. & Yap, A. S. (2013) Patterns in space: coordinating adhesion and actomyosin contractility at E-cadherin junctions, Cell Commun Adhes. 20, 201–12.

26. Varelas, X., Samavarchi-Tehrani, P., Narimatsu, M., Weiss, A., Cockburn, K., Larsen, B. G., Rossant, J. & Wrana, J. L. (2010) The Crumbs complex couples cell density sensing to Hippo-dependent control of the TGF-β-SMAD pathway, Dev Cell. 19, 831–44.

27. Nishioka, N., Inoue, K., Adachi, K., Kiyonari, H., Ota, M., Ralston, A., Yabuta, N., Hirahara, S., Stephenson, R. O., Ogonuki, N., Makita, R., Kurihara, H., Morin-Kensicki, E. M., Nojima, H., Rossant, J., Nakao, K., Niwa, H. & Sasaki, H. (2009) The Hippo signaling pathway components Lats and Yap pattern Tead4 activity to distinguish mouse trophectoderm from inner cell mass, Dev Cell. 16, 398–410.

28. Benham-Pyle, B. W., Pruitt, B. L. & Nelson, W. J. (2015) Cell adhesion. Mechanical strain induces E-cadherin-dependent Yap1 and β-catenin activation to drive cell cycle entry, Science. 348, 1024–7.

29. Acharya, B. R., Wu, S. K., Lieu, Z. Z., Parton, R. G., Grill, S. W., Bershadsky, A. D., Gomez, G. A. & Yap, A. S. (2017) Mammalian Diaphanous 1 Mediates a Pathway for E-cadherin to Stabilize Epithelial Barriers through Junctional Contractility, Cell Rep. 18, 2854–2867.

30. Citi, S. (2019) The mechanobiology of tight junctions, Biophys Rev. 11, 783–793.

31. Matter, K. & Balda, M. S. (2003) Signalling to and from tight junctions, Nat Rev Mol Cell Biol. 4, 225–36.

32. Zihni, C., Mills, C., Matter, K. & Balda, M. S. (2016) Tight junctions: from simple barriers to multifunctional molecular gates, Nat Rev Mol Cell Biol. 17, 564–80.

33. Zhao, B., Li, L., Lu, Q., Wang, L. H., Liu, C. Y., Lei, Q. & Guan, K. L. (2011) Angiomotin is a novel Hippo pathway component that inhibits YAP oncoprotein, Genes Dev. 25, 51–63.

34. Oka, T., Remue, E., Meerschaert, K., Vanloo, B., Boucherie, C., Gfeller, D., Bader, G. D., Sidhu, S. S., Vandekerckhove, J., Gettemans, J. & Sudol, M. (2010) Functional complexes between YAP2 and ZO-2 are PDZ domain-dependent, and regulate YAP2 nuclear localization and signalling, Biochem J. 432, 461–72.

35. González-González, L., Gallego-Gutiérrez, H., Martin-Tapia, D., Avelino-Cruz, J. E., Hernández-Guzmán, C., Rangel-Guerrero, S. I., Alvarez-Salas, L. M., Garay, E., Chávez-Munguía, B., Gutiérrez-Ruiz, M. C., Hernández-Melchor, D., López-Bayghen, E. & González-Mariscal, L. (2022) ZO-2 favors Hippo signaling, and its re-expression in the steatotic liver by AMPK restores junctional sealing, Tissue Barriers. 10, 1994351.

36. Xu, J., Kausalya, P. J., Van Hul, N., Caldez, M. J., Xu, S., Ong, A. G. M., Woo, W. L., Mohamed Ali, S., Kaldis, P. & Hunziker, W. (2021) Protective Functions of ZO-2/Tjp2 Expressed in Hepatocytes and Cholangiocytes Against Liver Injury and Cholestasis, Gastroenterology. 160, 2103–2118.

37. Spadaro, D., Tapia, R., Jond, L., Sudol, M., Fanning, A. S. & Citi, S. (2014) ZO proteins redundantly regulate the transcription factor DbpA/ZONAB, J Biol Chem. 289, 22500–11.

38. Domínguez-Calderón, A., Ávila-Flores, A., Ponce, A., López-Bayghen, E., Calderón-Salinas, J. V., Luis Reyes, J., Chávez-Munguía, B., Segovia, J., Angulo, C., Ramírez, L., Gallego-Gutiérrez, H., Alarcón, L., Martín-Tapia, D., Bautista-García, P. & González-Mariscal, L. (2016) ZO-2 silencing induces renal hypertrophy through a cell cycle mechanism and the activation of YAP and the mTOR pathway, Mol Biol Cell. 27, 1581–95.

39. Mangold, S., Wu, S. K., Norwood, S. J., Collins, B. M., Hamilton, N. A., Thorn, P. & Yap, A. S. (2011) Hepatocyte growth factor acutely perturbs actin filament anchorage at the epithelial zonula adherens, Curr Biol. 21, 503–7.

40. Gomez, G. A., McLachlan, R. W., Wu, S. K., Caldwell, B. J., Moussa, E., Verma, S., Bastiani, M., Priya, R., Parton, R. G., Gaus, K., Sap, J. & Yap, A. S. (2015) An RPTPalpha/Src family kinase/Rap1 signaling module recruits myosin IIB to support contractile tension at apical E-cadherin junctions, Mol Biol Cell. 26, 1249–62.

41. Leerberg, J. M., Gomez, G. A., Verma, S., Moussa, E. J., Wu, S. K., Priya, R., Hoffman, B. D., Grashoff, C., Schwartz, M. A. & Yap, A. S. (2014) Tension-sensitive actin assembly supports contractility at the epithelial zonula adherens, Curr Biol. 24, 1689–99.

42. Xu, J., Kausalya, P. J., Ong, A. G. M., Goh, C. M. F., Mohamed Ali, S. & Hunziker, W. (2022) ZO-2/Tjp2 suppresses Yap and Wwtr1/Taz-mediated hepatocyte to cholangiocyte transdifferentiation in the mouse liver, NPJ Regen Med. 7, 55.

43. Smutny, M., Wu, S. K., Gomez, G. A., Mangold, S., Yap, A. S. & Hamilton, N. A. (2011) Multicomponent analysis of junctional movements regulated by myosin II isoforms at the epithelial zonula adherens, PLoS One. 6, e22458.

44. Wu, S. K., Ariffin, J., Tay, S. C. & Picone, R. (2023) The variant senescence-associated secretory phenotype induced by centrosome amplification constitutes a pathway that activates hypoxia-inducible factor-1alpha, Aging Cell. 22, e13766.

45. Wu, S. K., Gomez, G. A., Michael, M., Verma, S., Cox, H. L., Lefevre, J. G., Parton, R. G., Hamilton, N. A., Neufeld, Z. & Yap, A. S. (2014) Cortical F-actin stabilization generates apical-lateral patterns of junctional contractility that integrate cells into epithelia, Nat Cell Biol. 16, 167–78.

46. Greenlees, R., Mihelec, M., Yousoof, S., Speidel, D., Wu, S. K., Rinkwitz, S., Prokudin, I., Perveen, R., Cheng, A., Ma, A., Nash, B., Gillespie, R., Loebel, D. A., Clayton-Smith, J., Lloyd, I. C., Grigg, J. R., Tam, P. P., Yap, A. S., Becker, T. S., Black, G. C., Semina, E. & Jamieson, R. V. (2015) Mutations in SIPA1L3 cause eye defects through disruption of cell polarity and cytoskeleton organization, Hum Mol Genet. 24, 5789–804.

47. Wu, S. K., Budnar, S., Yap, A. S. & Gomez, G. A. (2014) Pulsatile contractility of actomyosin networks organizes the cellular cortex at lateral cadherin junctions, Eur J Cell Biol. 93, 396–404.

48. Li, W., Cooper, J., Zhou, L., Yang, C., Erdjument-Bromage, H., Zagzag, D., Snuderl, M., Ladanyi, M., Hanemann, C. O., Zhou, P., Karajannis, M. A. & Giancotti, F. G. (2014) Merlin/NF2 loss-driven tumorigenesis linked to CRL4(DCAF1)-mediated inhibition of the hippo pathway kinases Lats1 and 2 in the nucleus, Cancer Cell. 26, 48–60.

49. Pan, M., Chew, T. W., Wong, D. C. P., Xiao, J., Ong, H. T., Chin, J. F. L. & Low, B. C. (2020) BNIP-2 retards breast cancer cell migration by coupling microtubule-mediated GEF-H1 and RhoA activation, Science Advances. 6, eaaz1534.

50. Han, S. P., Gambin, Y., Gomez, G. A., Verma, S., Giles, N., Michael, M., Wu, S. K., Guo, Z., Johnston, W., Sierecki, E., Parton, R. G., Alexandrov, K. & Yap, A. S. (2014) Cortactin scaffolds Arp2/3 and WAVE2 at the epithelial zonula adherens, J Biol Chem. 289, 7764–75.

51. Wu, S. K. & Priya, R. (2019) Spatio-Temporal Regulation of RhoGTPases Signaling by Myosin II, Front Cell Dev Biol. 7, 90.

52. Deng, L., Ho, C., Picone, R., Abderazzaq, F., Flanagan, N., Low, B. C. & Wu, S. K. (2024) The Senescence-Associated-Secretory Phenotype constituting HIF-1α activation is independent of micronuclei induction and associated cGAS-mediated interferon response, bioRxiv, 2024.09.25.615081.

53. Pan, M., Chew, T. W., Wong, D. C. P., Xiao, J., Ong, H. T., Chin, J. F. L. & Low, B. C. (2020) BNIP-2 retards breast cancer cell migration by coupling microtubule-mediated GEF-H1 and RhoA activation, Sci Adv. 6, eaaz1534.

54. Wu, S. K., Gomez, G. A., Stehbens, S., Acharya, B. R., Ratheesh, A., Priya, R., Lagendijk, A. & Bershadsky, A. (2022) Editorial: Forces in biology - Cell and developmental mechanobiology and its implications in disease - Volume II, Front Cell Dev Biol. 10, 1082857.

55. Wu, S. K., Gomez, G. A., Stehbens, S. J. & Smutny, M. (2020) Editorial: Forces in Biology - Cell and Developmental Mechanobiology and Its Implications in Disease, Front Cell Dev Biol. 8, 598179.

56. Wu, S. K., Ho, C. Z., Sun, F., Lou, Y., Huang, C. B.-X., Xiao, J., Shagirov, M., Yow, I., Chin, J. F. L., Verma, S., Yap, A. S., Lin, Y., Hiraiwa, T. & Low, B. C. (2023) A morphogenic cascade emerges from the co-evolution of spheroid fluidization and fracturing of multicellular barriers, bioRxiv, 2023.09.25.559247.s

